# The rate of inversion fixation in plant genomes is highly variable

**DOI:** 10.1101/2022.08.31.506062

**Authors:** Kaede Hirabayashi, Gregory L. Owens

**Affiliations:** Department of Biology, University of Victoria, Victoria, BC, Canada

## Abstract

Chromosomal inversions are theorized to play an important role in adaptation by preventing recombination, but testing this hypothesis requires an understanding of the rate of inversion fixation. Here we use chromosome-level whole genome assemblies for 32 genera of plants to ask how fast inversions accumulate and what factors affect this rate. We find that on average species accumulate 4 to 28 inversions per million generations, but this rate is highly variable, and we find no correlation between sequence divergence or repeat content and the number of inversions and only a small correlation with chromosome size. We also find that inversion regions are depleted for genes and enriched for TEs compared to the genomic background. This suggests that idiosyncratic forces, like natural selection and demography, are controlling how fast inversions fix.

## Introduction

The field of genomics has undergone a remarkable expansion in the last decade. With the rapid advances in long- and linked-read sequencing technologies, assembling a chromosome-resolved plant genome is no longer a fantasy (Pucker et al. 2022). While earlier draft genomes often covered the entire genome, they were in many small contigs which meant that genome structure was not resolved. Recent empirical work has highlighted that changes in genome structure can be critical for important evolutionary processes such as adaptation and speciation, and chromosome-resolved genome assemblies allow for these to be surveyed in an unbiased way for the first time (Mérot et al. 2020).

One type of structural variation, inversions, are particularly interesting because of their effect on meiosis and recombination. Initial investigations into inversions focused on underdominance effects (White 1973). Due to how homologous alignment occurs during meiosis, in heterozygous individuals a recombination event within an inversion will lead to unbalanced gametes and a loss of fertility (Dobzhansky 1933; White 1978). This underdominance led to research into their role in reproductive isolation, as inversions in different orientation are often fixed between species (Trickett and Butlin 1994; Rieseberg 2001) and have been shown to directly cause hybrid sterility (reviewed in L. Zhang, Reifová, Halenková, & Gompert, 2021). It was also recognized that the recombination suppression abilities of inversions may link together favourable alleles and be relevant in the context of adaptation (Sturtevant and Mather 1938; Ohta and Kojima 1968; Charlesworth and Charlesworth 1973; Kirkpatrick and Barton 2006). For example, loci that are locally adaptive can benefit from inversions since they can be co-inherited as an adaptive multi-loci haplotype and protected from recombination with non-adaptive alleles (Kirkpatrick and Barton 2006). Numerous non-model systems have been observed to have inversions with adaptive significance: sunflower (Todesco et al. 2020), monkeyflower (Lowry and Willis 2010), rainbow trout (Leitwein et al. 2017), Atlantic cod (Barth et al. 2017; Sodeland et al. 2022), stickleback (Jones et al. 2012), deer mice (Hager et al. 2022; Harringmeyer and Hoekstra 2022).

There are several molecular mechanisms of inversions, all of which are triggered by some form of DNA strand breaks during meiosis or in other situations (reviewed in detail by Burssed, Zamariolli, Bellucco, & Melaragno, 2022; Casals & Navarro, 2007). In non-allelic homologous recombination (NAHR), segments with a high degree of sequence similarity misalign during meiosis and recombine within the chromatid (intrachromatid) instead of between sister chromatids. This is known as ectopic recombination. When repeats are found in the opposite orientation, inversion of the region surrounded by the misaligned repeats can occur. This means that inverted repeats are necessary for inversions mutations with this mechanism. Non-homologous end-joining (NHEJ) inversions occur from random paired double-stranded breaks surrounding a segment of the genome, followed by 180° flipping of the detached region and a repair. This mechanism does not require any sequence homology around breakpoints and could occur during any stage of the cell cycle. The isochromatid mechanism is similar to NHEJ but requires staggered single-stranded breaks instead of double-stranded breaks (Ranz et al. 2007). The inversion through this mechanism would result in the creation of inverted sequence duplicates around the breakpoint regions. Lastly, inversions that are especially small and found in a complex rearrangement pattern could arise from DNA replication machinery being disturbed and confused due to formation of DNA secondary structures (e.g., Cruciform, hairpin, triple helical DNA, quadruplex). The replication fork may be subjected to breakage or stalling and cause template switching event, called invasion. If the new position has switched orientations of strand (so say invasion occurred from template to lagging strand), then inversions of that segment would result. Additionally, large number of mobile elements (transposable elements or TEs) has been associated with recurrent chromosomal rearrangements, documented by the reuse of some of the breakpoints identified (Ranz et al. 2007). These regions could be prone to inversions and be the driving force of rearrangements.

Despite growing interest of chromosomal inversions in adaptive potential and appreciation of their prevalence, there is a lack of knowledge on the rate of inversion fixation between species in plants. Most studies of the rate of chromosomal inversions have focused on *Drosophila* species (Ranz et al. 2001; Bhutkar et al. 2008). A literature review on plants has estimated the rate of inversion fixation to be around 15-30 inversions per million years based on the divergence time of a small number of species analyzed (Huang and Rieseberg 2020). However, the method of inversion detection and divergence estimation cited in this review were variable and did not account for the size of the inversions. Thus, further study with robust and consistent approach is needed.

Leveraging the resource for publicly available genome assemblies, we conducted a comparative study to investigate how chromosomal inversions accumulate in eudicots. We expected to observe that inversions would accumulate over time at a consistent rate; therefore, more divergent species pairs should harbour more inversions. We also predicted that if there is more space in the genome for chromosomal rearrangement (i.e., larger genome size), would lead to more inversions. Alternatively, if fixation of inversions was dominated by selection, then rates will be idiosyncratic. We further investigated the genomic context of inversions, which gives us insight into their functional significance. Lastly, we asked if inversions were surrounded by inverted repeats, which tells about the relative rate of different inversion mechanisms.

## Methodology

### Data collection

Publicly available genomes within Eudicots were searched through the published plant genomes database (last screened in November 2021; https://www.plabipd.de/plant_genomes_pa.ep). We selected genera with at least two sequenced species. Then individual genomes were screened for all of the following criteria: 1) same ploidy (diploid), chromosome-resolved assembly, 3) same number of chromosomes, 4) *de novo* assembled and 5) valid NCBI BioProject number or similar. Assemblies published before 2018 were carefully assessed for quality. Those that used guidance from genetic mapping or reference assembly were excluded. If the assemblies did not have an associated publication or provided no adequate information on how exactly the genome was sequenced, they were excluded from the study. For one species per genus, we downloaded coding sequences and gene annotations. In total, 64 chromosome-resolved genomes from 32 genera were included (Supplementary Table 1). To explore biological factors that may affect the fixation of inversions, we surveyed the literature to identify whether the species have evidence of hybridization and their reproductive self-compatibility. Evidence of hybridization was categorized into three groups: weak (no evidence of hybridization or evidence of sterile hybrid attempt), strong (evidence of hybridization/introgression in nature or lab or genes), unclear (lack of evidence to support either weak or strong). Self-compatibility was assessed by three categories: selfing (mainly reproduce by self-pollinating), mixed (can self or mate), self-incompatible (cannot produce viable offspring from selfing). Two genome pairs were missing the gene annotation file and thus excluded from analyses involving gene locations. All species used and recorded information are in Supplementary Table 1.

### Structural analysis

The collected 32 genome pairs were analyzed for structural variant detection using Synteny and Rearrangement Identifier v1.5 (SyRI) (Goel et al. 2019). To do this, first, genomes were assigned either reference or query based on assembly statistics (N50 and number of contigs) and availability of gene annotation file. ‘Reference’ status was assigned to those with good contiguity and gene annotation file available. Next, only chromosomal sequences were extracted from genome assemblies and scaffolds were removed. Therefore, from here onward, the ‘total genome length’ in our study refers to the total length of assembly that has been assigned to chromosomes. For *Luffa* and *Acer* genomes, the chromosomes in the query genome that had half synteny with one chromosome and the other half with another chromosome in the reference genome were removed from SyRI analysis. This is due to SyRI being unable to perform comparison between divergent genomes lacking one-to-one synteny. Specifically, in *Luffa*: CM029395, CM029402 (reference), CM022716, CM022722 (query) were excluded, in *Acer*: chr3, chr4 (reference), CM017761, CM017766 (query) were excluded. When running SyRI, default parameters were used and whole-genome base-to-base alignment was performed by minimap2 v2.17-r974-dirty (Li 2018, 2021). When necessary, chromosomes were reverse complemented using Samtools v1.12 (Li et al. 2009) prior to SyRI.

For our analyses, we identified inversions that differ in orientation between closely related species, but we are not able to determine which orientation is derived or ancestral. An inversion region in the reference species is a region that is in the opposite orientation in the query species, which is because either the reference or query species inverted. If we assume that the rate of inversion fixation is roughly equal between the two species – which may not be true – then roughly half of the inversion regions contain a derived inversion in the reference species and the other half contain the ancestral state, but have inverted in the query species.

### Transposable element annotation

Transposable elements (TEs) were detected and annotated using Extensive De novo TE Annotator v2.0.0 (EDTA) (Ou et al. 2019) pipeline on the 30 reference genomes with gene annotations available. Due to resource limitation, the pipeline was performed by chromosome-by-chromosome. The whole genome fasta file was first divided into individual chromosome files. Then the EDTA pipeline was performed separately on each chromosome as follows and the resulting TE library was later combined. In brief, candidate TE sequences were *de novo* identified using LTR-Finder (Xu and Wang 2007; Ou and Jiang 2019), LTRharvest (Ellinghaus et al. 2020), LTR_retriever (Ou and Jiang 2018), generic repeat finder (Shi and Liang 2019), and HelitronScanner (Xiong et al. 2014) respectively. Once individual TEs were identified, candidates were filtered (Zhang et al. 2019) and further refined by RepeatModeler (Flynn et al. 2020) according to EDTA pipeline default parameters. Finally, TE-free coding sequences retrieved from gene annotation files (feature name = ‘CDS’) were aligned to the repeat library, and those overlapped with coding sequence (CDS) were excluded from the identified TE candidates. Total genomic TE content (%) was calculated by the total length of detected TE divided by the total genome length.

### Species divergence calculation

The principal advantage of SyRI over basic genome aligners is that SyRI identifies larger regions with consistent synteny. To do this, SyRI takes alignment blocks identified by the initial aligner, in this case minimap2, and identifies larger regions consisting of multiple consecutive one-to-one alignment blocks. We focused our analysis on regions which each represent a single structural variant (or contiguous syntenic region without any structural variation). While SyRI identifies several types of structural variation, we focused our analysis on inversions and syntenic regions only. To minimize noise from small amounts of data, detected regions smaller than 1 kbp were excluded from the dataset. To calculate the sequence divergence for each region (*X*) we used equation 1. From SyRI output, for each block (*i*) in a region, we measured percent identity (*x*_*i*_) and reference length (*L*_*i*_).

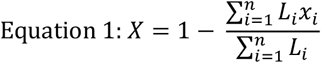

To calculate the average sequence identity between species, we used the equation 1 but, in this case, combining all syntenic blocks. With this formula, sequence divergence score of 0 indicates identical species, and the value increases as the genomes become more different.

To convert sequence divergence into divergence time we used the genome-wide substitution rate estimated from *Arabidopsis thaliana* (Exposito-Alonso et al. 2018), which estimated 2 to 5 × 10^− 9^ substitutions per year. We note that *Arabidopsis* is an annual plant, while the species studied here have varying generation times which may affect substitution rates, but estimates of mutation rates from somatic tissue of plants found a range of values which encompassed the substitution rate used here (Wang et al. 2019). Based on this rate, and that the divergence rate is equal for each species in the pair, each percentage point divergence represents 1 to 2.5 million years of separation. This estimate should be used with caution as substitution rates are known to vary and divergence between close relatives is affected by standing variation (Ho et al. 2011). We present our rates of inversion accumulation in years of evolution. This means that a species pair that had a common ancestor 1 million years ago, is separated by 2 million years of evolution accounting for evolution in both species.

### Quantifying cds and te in inverted regions vs. syntenic regions

We were interested if inversions contained different proportions of genomic elements. Bedtools v2.30.0 (Quinlan and Hall 2010) intersect function was used to analyze how many features and base pairs of coding sequence (CDS; as identified by annotations) or TEs (as identified by EDTA) were found to overlap with the inverted regions in the genome. Inversion regions identified by SyRI were extracted using custom perl script and reorganized into a bed format using Linux commands. The gene annotation file and TE library in gff3 were reformatted into bed format using gff2bed. Prior to using bedtools intersect, CDS and TE bed files were edited by bedtools merge function to concatenate overlapping features into a single feature, avoiding overrepresentation due to isoforms of the same gene sometimes present in the gene annotation file. The validity of data was confirmed by ensuring the proportion of CDS or TE per inverted region no larger than 1. In addition to inverted regions, the process was repeated with syntenic regions identified by SyRI.

### Quantifying CDS and TE in inversion breakpoint regions

We were also interested in whether genomic elements differ at the breakpoints of inversions compared to the rest of the genome. Quantification of CDS/TE in inversion breakpoints were performed using Bedtools (Quinlan and Hall 2010) intersect function. The number of CDS and TE in the inversion breakpoint regions were extracted as follows. First, inversion breakpoint regions (defined as 4 kbp regions downstream or upstream of breakpoints start or end respectively) were identified. A small number of inversion breakpoint regions were within 4 kbp of the end of the chromosome, and were excluded from the dataset. A total of 11,225 breakpoint regions from 30 genus pairs were used for analysis. To infer whether breakpoints are enriched with CDS/TE, baseline numbers for the whole genome are necessary. To determine the baseline number of CDS/TE in the genome, the total length of CDS/TE was computed from the merged bed file from previous step. Then the genomic CDS/TE proportion was calculated by the total length of CDS/TE divided by the length of the reference genome. The resulting average proportion of CDS/TE in the 4 kbp breakpoint regions was compared to the genomic CDS/TE proportion.

### Quantifying frequency of gene occurrence at inversion breakpoints

Chromosomal inversions can disrupt gene sequence if an inversion breakpoint occurs within the gene itself. Since we cannot identify whether the derived orientation occurred in the reference or query genome, we are focusing on genes identified in the reference species. If an inversion breakpoint falls within a gene sequence it has two possible meanings (Figure 1):

**Figure 1:**
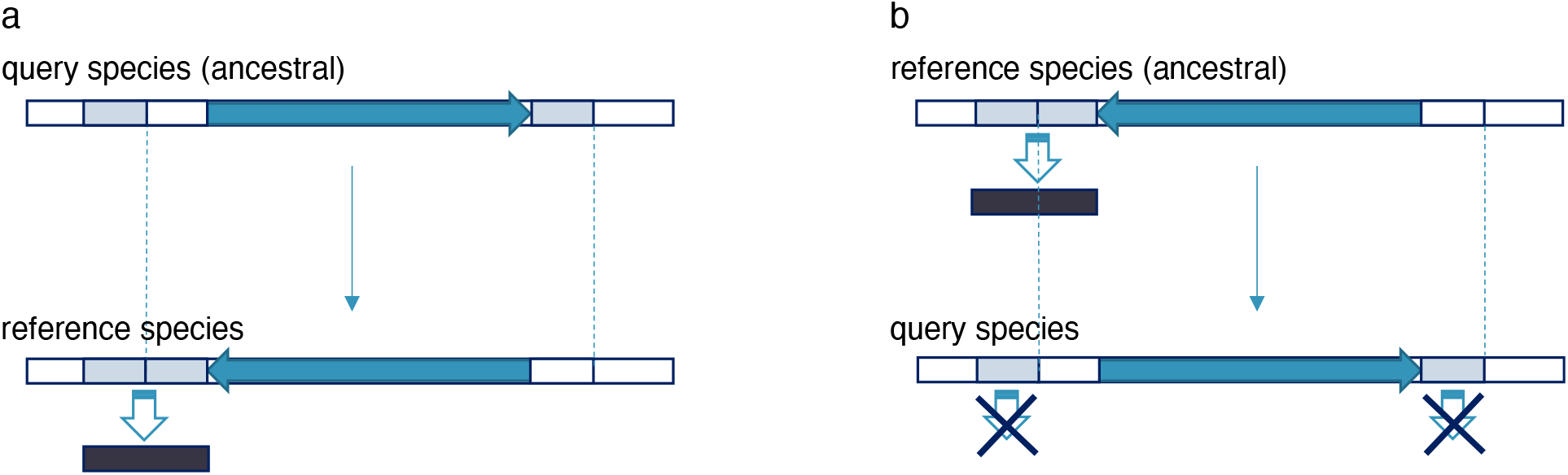
Two different interpretations of inversion consequences. **a**) An inversion creates a new gene in the reference species. **b**) An inversion disrupts an ancestral gene in the query species.

1. The orientation is derived in the reference species. The inversion either created the gene or modified its coding sequence.
2. The orientation is ancestral in the reference species. The inversion disrupted or modified the coding sequence of the gene in the query species.

Both models assume that the genes are largely shared between the reference and query species. For example, if a new gene appears only in the reference species genome it cannot be disrupted by an inversion in the query genome.

We were interested in how often inversion breakpoints fell within genes. We used Bedtools intersect to count the number of breakpoints that occurred within the coding sequence or introns of a gene. To identify if breakpoints are less likely to occur within genes, we selected *n* random positions (*n* adjusted to 1000 per chromosome for each genome) from the syntenic portion of the genome using Bedtools (Quinlan and Hall 2010) random function and used this as a baseline. Additionally, we also calculated the proportion of genes that contained at least one inversion breakpoint to identify how often inversions may be disrupting genes.

### Breakpoints regions sequence similarity analysis

Some models for inversion mutations require or create segmental duplications at the breakpoints of the inversion. To explore this idea, the 10 kbp regions upstream and downstream of breakpoints described above were aligned to each other using BLAST v2.12.0 (Zhang et al. 2000). In some cases, breakpoints were within 10 kbp of the ends of chromosomes, resulting in smaller analysis regions. For these, we required there to be at least 2 kbp of sequence both upstream and downstream of the inversion. In cases where one reference sequence (breakpoint start region) had multiple BLAST hits to the query sequence (breakpoint end region), only the longest aligned hit was retained.

## Results

### Inversion accumulation is idiosyncratic

We identified a total of 6,140 inversions across our 32 comparisons, 5,298 of which were larger than 1000 bp. In general, inversions tended to be small: 45.0% (2,766/6,140) of inversions were less than 10 kbp, 46.8% (2,873/6,140) were between 10 kbp and 1 Mbp and 8.2% (501/6,140) were greater than 1 Mbp (Figure 2a). The number of inversions varied between comparisons (Figure 2b), but in general represented a relatively small proportion of the total genome length, from 1.3% to 37.4% (Figure 2c). For each of our genera, we observed whether sequence identity in inversion regions differed from sequence identity in syntenic regions and found no significant difference for all 32 comparisons (Supplementary Figure 1). In syntenic regions, our species had on average 3% to 5% sequence divergence, supporting their close relationship (Supplementary Figure 2).

**Figure 2:**
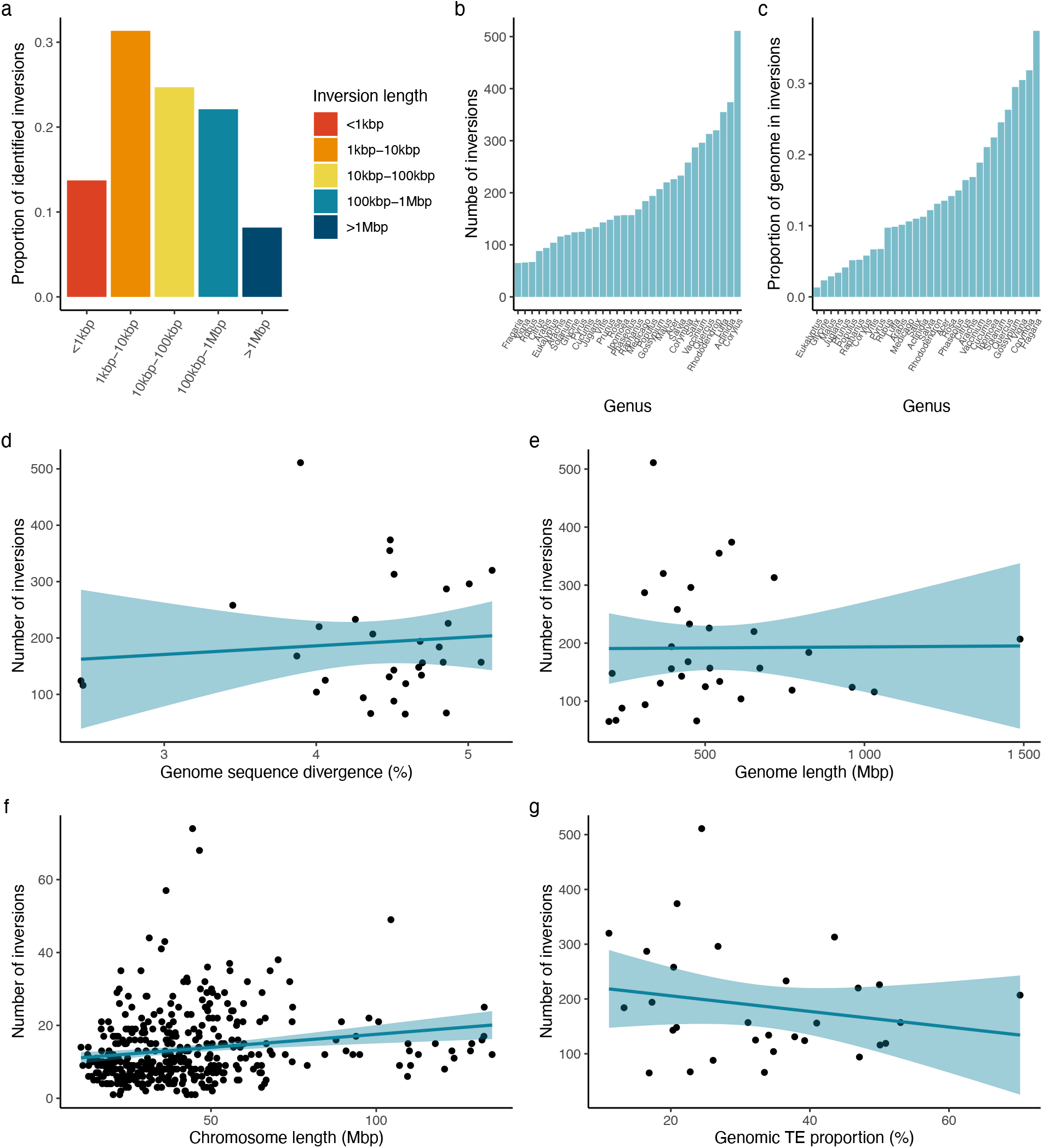
Summary of inversions between paired eudicot genomes in the same genus. **a**) Size distribution of inversions by inversion length. **b**) Number of inversions and **c**) proportion of reference genome in inversion, for 32 species pairs used in the study. Number of inversions plotted against **d**) percent sequence divergence between the 32 species pair (F value = 0.264, *p* = 0.611), **e**) genome length of the reference species genome (n = 32; F value = 0.002, *p* = 0.961), **g**) genomic TE proportion calculated by total length of TE / reference genome length (n = 30; F value = 1.139, *p* = 0.295). Each datapoint represents a paired species within the same genus. **f**) Number of inversions per chromosome plotted against its length (n = 405; F value = 12.78; *p* = 0.00039 ***; linear model R^2^ = 0.029).

We were interested in determining factors that affected the number of inversions and the rate of inversion accumulation. Sequence divergence is expected to increase over time and can be a proxy for divergence time (with many potential caveats) (Drummond et al. 2006; Ho et al. 2011). We used a linear model to ask if species with higher sequence divergence had more inversions and found no significant relationship (*p* = 0.6) (Figure 2d). This was also not significant if we instead used the proportion of the genome that is inverted, rather than the number of inversions (*p* = 0.5) (Supplementary Figure 3). All else being equal, we expected that larger genomes should have more inversions. When tested using the entire genome length, we saw no significant relationship (*p* = 0.96) (Figure 2e), but when treating each chromosome separately we found a slight positive relationship (n = 405; F value = 12.78; *p* = 0.00039 ***; linear model R^2^= 0.029) (Figure 2f). Since inversion origin is often associated with DNA repeats, we expected that species with a higher proportion of TEs would have more inversions, but again this relationship was not significant (*p* = 0.3) (Figure 2g).

Given that inversions are not accumulating in a clock-like manner, we used a one-way ANOVA to explore other factors controlling the number of inversions. We found that the assembly method used, sequencing technology and whether the species had evidence of hybridization did not affect the number of inversions significantly (Table 1), but reproductive strategy did when it was calculated with the proportion of inverted genome instead of the number (*p* = 0.031 *). We found that selfing species had the highest proportion of genome inverted, while mixed and self-incompatible species had a similar proportion (Supplementary Figure 4).

**Table 1.** One-way ANOVA results. Tests performed for the number of inversions and proportion of reference genome in inverted orientation against the following five factors: evidence of hybridization (weak/strong), reproductive strategy (selfing/mixed/self-incompatible), genome assembler category (accurate/fast or not resource intensive/short-read based), long-read sequencing platform (Oxford Nanopore/Pacific Bioscience/none or short-read only), whether physical mapping (i.e., Hi-C, BioNano optical map, long-range Chicago) is performed or not. The reproductive strategy and assembly methods were assigned to each genome assembly (reference and query) separately, then the paired category was used in the stats (n = 32).

| Inversion measure | Number of inversions |  | Proportion of genome inverted |  |
| --- | --- | --- | --- | --- |
|  | F value | Pr(>F) | F value | Pr(>F) |
| Evidence of hybridization | 0.317 | 0.578 | 0.001 | 0.973 |
| Reproductive strategy | 1.73 | 0.163 | 2.942 | 0.031 * |
| Assembler category (accurate, fast, short-read based) | 1.223 | 0.324 | 0.73 | 0.579 |
| Sequencing platform (ONT vs PacBio vs short-read only) | 0.772 | 0.52 | 1.504 | 0.235 |
| Physical mapping (Y/N) | 1.679 | 0.204 | 0.023 | 0.977 |

\* STATISTICALLY SIGNIFICANT (P < 0.05)

Although the rate of inversion accumulation relative to sequence divergence is not consistent in our dataset, the range of possible values may be a useful baseline for other studies. We present a fixation rate by dividing the number of inversions by the sequence divergence for different size categories of inversions (Figure 3). We find the smaller inversions are more common, with the exception of the smallest category. Further, based on an estimate of 1 to 2.5 million years of divergence per 1% sequence divergence, we present a range of estimates of inversion accumulation by size (Table 2).

**Figure 3:**
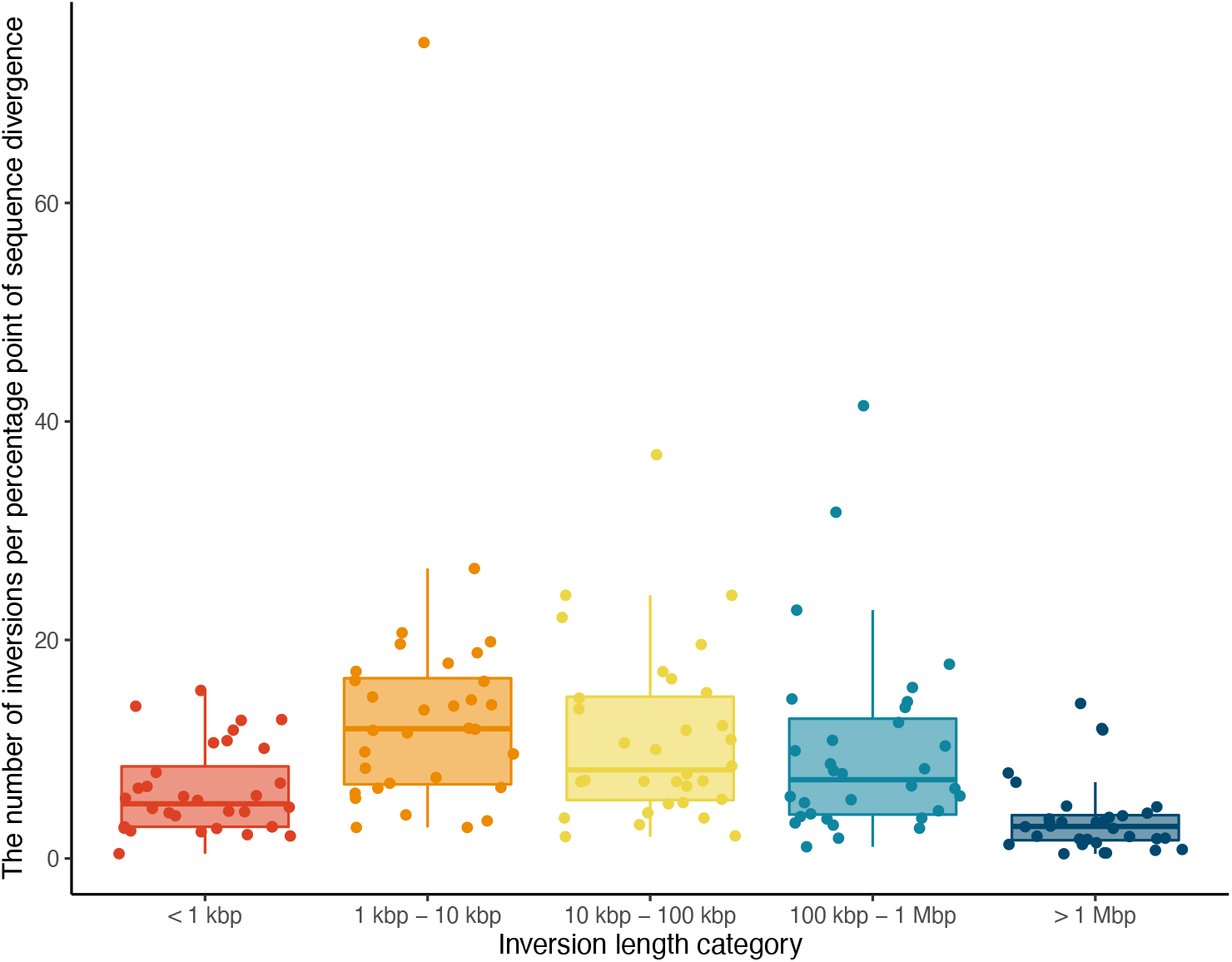
Rate of inversion fixation by inversion size. Rate of inversion occurrence calculated as number of inversions per percentage point of sequence divergence by different size categories (n = 32 for each category).

**Table 2.** Rate of inversion accumulation by inversion length with respect to species divergence time. Species divergence time was estimated using Arabidopsis *thaliana* substitution rate (Exposito-Alonso et al. 2018).

| Inversion length category | Mean number of inversions per 1% seq divergence (25<sup>th</sup> – 50<sup>th</sup> – 75<sup>th</sup> percentile) |  |  | Mean number of inversions per estimated species divergence time (million years ago, 25<sup>th</sup> – 50<sup>th</sup> – 75<sup>th</sup> percentile) |  |  |  |  |  |
| --- | --- | --- | --- | --- | --- | --- | --- | --- | --- |
| Conversion |  |  |  | 1 mya / 1% seq divergence |  |  | 2.5 mya / 1% seq divergence |  |  |
| < 1 kbp | 2.9 | **5.0** | 8.4 | 1.5 | **2.5** | 4.2 | 0.6 | **1.0** | 1.7 |
| 1 kbp – 10 kbp | 6.8 | **11.9** | 16.5 | 3.4 | **6.0** | 8.4 | 1.4 | **2.4** | 3.3 |
| 10 kbp – 100 kbp | 5.3 | **8.1** | 14.8 | 2.7 | **4.0** | 7.4 | 1.1 | **1.6** | 3.0 |
| 100 kbp – 1 Mbp | 4.0 | **7.2** | 12.8 | 2.0 | **3.6** | 6.4 | 0.8 | **1.4** | 2.6 |
| > 1Mbp | 1.7 | **2.9** | 4.0 | 0.8 | **1.5** | 3.0 | 0.3 | **0.6** | 0.8 |
| All | 20.7 | **35.1** | 56.5 | 10.3 | **17.5** | 28.2 | 4.1 | **7.0** | 11.3 |

### Inversions tend to occur in less functional regions

If inversions cause the deleterious disruption of gene sequence or expression, then we expect fixed inversions to contain less gene sequence when compared to syntenic regions. Indeed, we found this was the case (Figure 4a; t = −13.97, df = 29, *p* = 2.08e-14 ***). We also found the reverse pattern for TEs, where inversions were enriched compared to syntenic regions (Figure 4b; t = −3.93, df = 29, *p* = 4.77e-4 ***).

**Figure 4:**
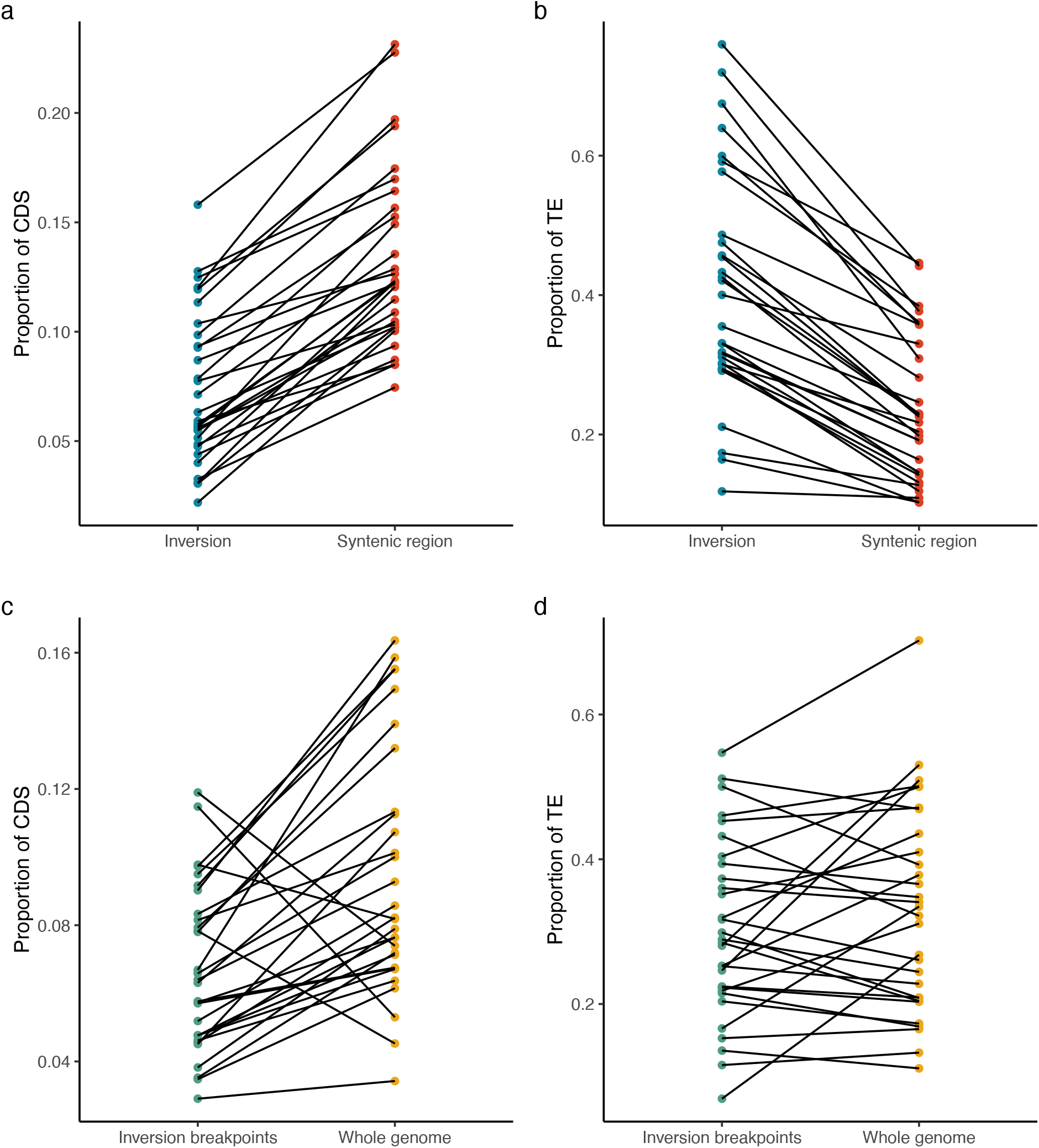
Genomic context of inversions and surrounding regions. The mean proportion of **a**) coding sequence and **b**) transposable elements inside inversion (blue) or syntenic region (red) (n = 30; *p* = 2.08e-14***, 4.77e-4***, respectively). The mean proportion of **c**) coding sequence and **d**) transposable elements in 4 kbp breakpoint regions (green) compared to the genomic CDS and TE proportion (yellow) in the genome (n = 30; *p* = 0.000107 ***, 0.142, respectively). Each datapoint represents a genus pair and the same species pair is connected by the line.

We next asked whether inversion breakpoints differed from the genome background. Although SyRI identifies breakpoints, limitations in the alignment of repetitive regions means that these may not be the exact breakpoint, therefore we selected 4 kbp regions downstream/upstream of identified breakpoint positions. These regions were compared against the genomic CDS and TE content in reference genome. We found that there was a lower proportion of CDS in breakpoint regions (Figure 4c; *p* = 0.000107 ***), but no consistent trend for TEs (Figure 4d; *p* = 0.142).

To infer whether inversions are disrupting gene sequence, or alternatively involved in the creation of new genes, we counted the number inversion breakpoints (start/end) that fell within a gene (Figure 5a). Strikingly high occurrence of inversion breakpoints at a gene was observed in several genera (Figure 5a). Notably *Vigna, Ipomoea, Medicago, Solanum, Cucumis, Salix*, and *Phaseolus* had > 50% of breakpoints occurring within a gene. Despite this, the relatively small number of inversions compared to genes means that < 1% of genes contained an inversion breakpoint (Figure 5b).

**Figure 5:**
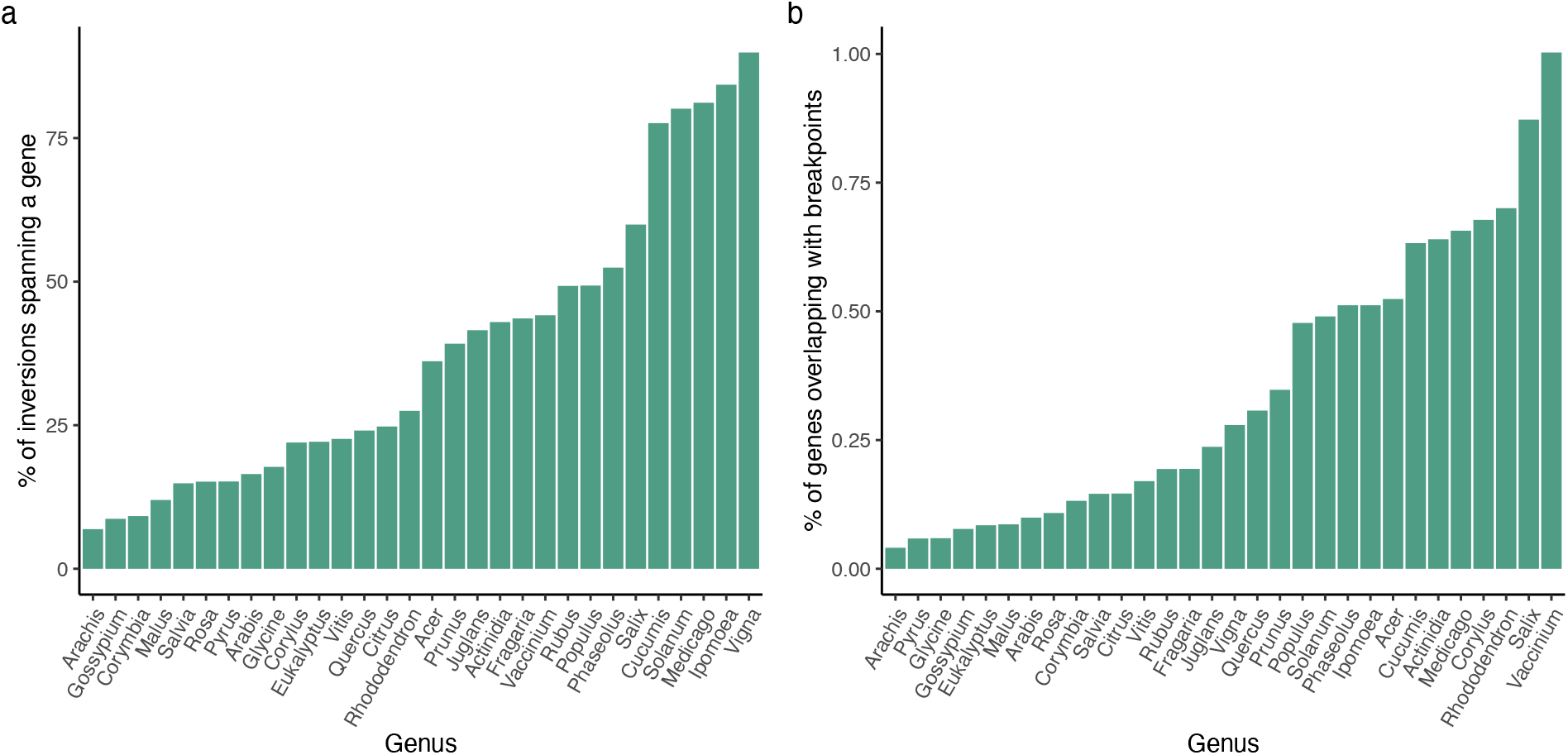
Inversion breakpoints and genes overlap. **a**) The percentage of inversion breakpoints that overlapped with a gene by genus. **b**) The percent of genes in reference genome that contained an inversion breakpoint.

### Inversions are most often not surrounded by repeats

Lastly, to test the molecular mechanism of inversions, we specifically looked for duplicates or inverted repeats surrounding the inversion breakpoints. Among the 5,626 tested 10 kbp breakpoint regions pair, 1,160 (20.6%) regions pair resulted in at least one BLAST hit. Of these, 592 were in the forward orientation and 568 were in inverted orientation (Figure 6).

**Figure 6:**
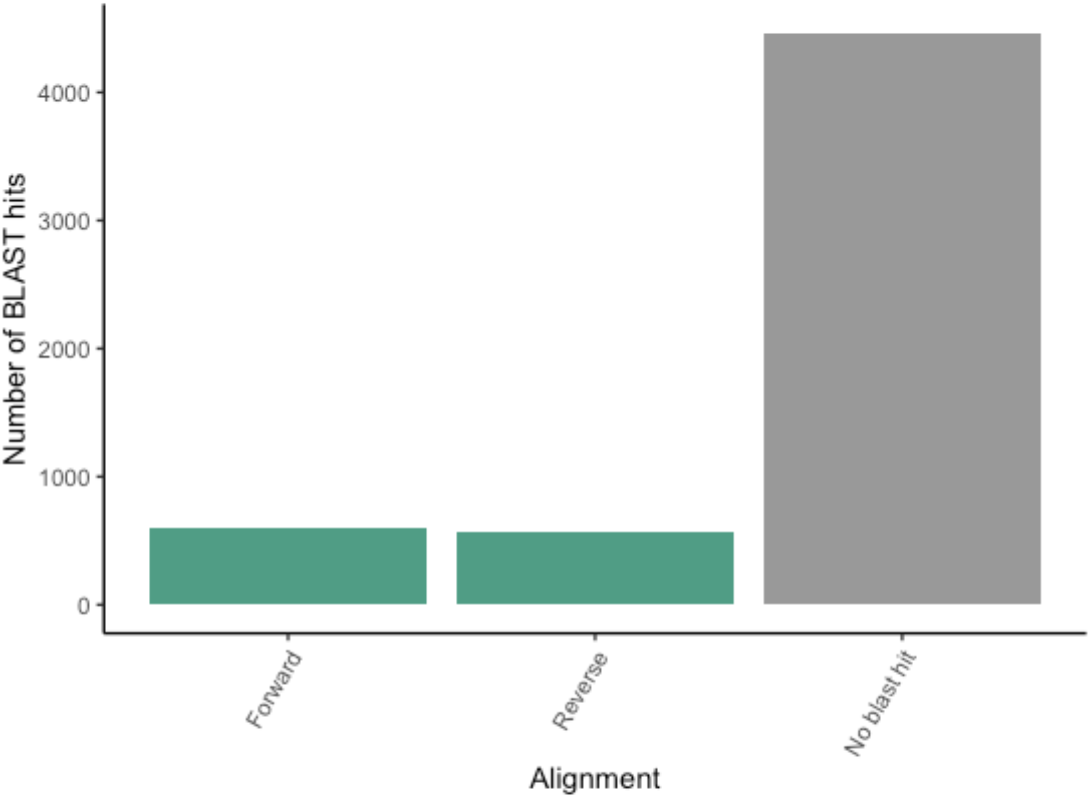
Rare presence of duplicates or inverted repeats near breakpoints. Summary of alignment between the 10 kbp window surrounding the 5,626 inversions detected using BLAST.

## Discussion

### The rate of inversion accumulation

Several high-profile studies have highlighted the adaptive importance of inversions. For example, the prairie sunflower (*Helianthus petiolaris*) has adapted to the dune environment while continuing to exchange genes with non-dune neighbouring populations (Huang et al. 2020). Recent studies have shown that the alleles controlling dune adaptation are found within large inversion regions suggesting the possibility that the recombination suppression of inversions is playing a role in maintaining adaptations (Huang et al. 2020; Todesco et al. 2020). One challenge to this hypothesis is that the number and size of inversions is not known so null models are challenging to parameterize. For example, if inversions are ubiquitous across the genome, we expect adaptive variation to be found in them regardless of other features of inversions.

To measure the rate of inversion fixation, we aligned chromosome-level genome assemblies for two species within the same genus. These comparisons explore how fast inversions fix in the genome by comparing the sequence divergence between species with the number of inversion differences. We are treating the rate as a baseline in comparison to systems like *H. petiolaris* where there is evidence that inversions are positively selected, but we recognize that the inversions themselves may not be neutral and that each species has its own demographic and adaptive history. Surprisingly, we find no correlation between sequence divergence, which is a proxy for coalescence time, and the number of inversions separating species (Figure 2d), suggesting that the rate of accumulation is dependent on factors not consistent between genera. Based on our data, the middle 50% of comparisons accumulated 20 to 57 inversions per 1% sequence divergence. Converting from sequence divergence to divergence time is fraught because it is affected by generation time, the mutation rate, and other factors which are likely to vary between our genera but it is helpful for scaling expectations (Ho et al. 2011; Wang et al. 2019). Based on substitution rates from *Arabidopsis*, we estimate 4 to 28 inversions per million years which is within the range estimated by Huang et al. from a more limited dataset (Huang and Rieseberg 2020).

Although we present a range of inversion accumulation rates, our primary finding is that this value is not consistent. There are several reasons — both technical and biological — that could explain the variation. Our inversion counts are based on whole genome alignments which is much better at capturing small inversions than genetic mapping but are not without error. Artifactual inversions can be introduced during assembly or scaffolding. We looked for the effect of sequencing and assembly method and failed to find a significant relationship, but our test is underpowered for the number of categories based on our sample size. There are also limitations in our ability to detect inversions based on alignments. We find that smaller inversions are more common, except for the smallest category (Figure 2a), which may reflect the limits of detectability or a trade-off between the high sensitivity to detect smaller inversions while minimizing the erroneous inverted alignments, especially in repetitive regions.

Different taxa have diverse genomic properties. Some may have inherently high recombination rate or epigenetic patterns that allow them to harbour chromosomal rearrangements more easily (Henderson 2012; Lloyd 2022). Each species also has its own demographic and selection history. Several of the species used in this analysis are domesticated crops, which have likely undergone greater bottlenecks and distinct selection pressures compared to wild species. This could lead to greater fixation of deleterious inversions similar to mutational load seen in some domestic species (Bosse et al. 2019).

The rate of inversion fixation is expected to be related to the rate of inversion mutation. Logically, longer genomes should have more opportunity for double-stranded breaks and therefore more inversion mutations. When considering each genome as a replicate we do not see a significant relationship, although there is a relatively small but significant relationship when treating chromosomes as replicates (Figure 2f). Higher repeat content is also often associated with chromosomal rearrangements, and for some mechanisms of inversion generation, repeats are necessary. We therefore expected that genomes with more transposable elements would have more inversions, but again we do not see a significant relationship (Figure 2g). We observed a clear linear relationship between the genome size and TE content (data not shown) as expected, but the occurrence of inversions seemed to be correlated with neither. Since neutral genomic features seem to play a relatively small role, we suggest that selection and demography are key in inversion fixation.

### Natural selection and inversions

A consistent rate of inversion accumulation makes an underlying assumption that the inversions themselves are neutral. If inversions are primarily fixed through positive selection, they likely represent a small fraction of total inversion mutations, and their count would be highly dependent on the distribution of fitness effects of new inversions. If inversions are instead primarily deleterious, then their fixation is dependent on the amount of genetic variation, the amount of selection on heterozygotes and homozygotes as well as the population structure of the species (Lande 1984). In both cases, the fixation rate is going to vary due to species specific effects (e.g., demographic history) as well as inversion specific effects (e.g., inversion selection coefficient) which lead to inconsistent fixation rates.

New inversions are expected to be underdominant because of disruptions to meiosis, and this should be stronger in larger inversions (White 1978; Rieseberg 2001). For primarily selfing species, this should be less of a barrier because of reduced heterozygosity (Hedrick 1981; Hoffmann and Rieseberg 2008). We see this play out in our ANOVA of mating system which showed the highest amount of the genome in inversions for comparisons involving selfing species (Supplementary Figure 4).

Inversions can play a role in speciation by containing reproductive isolation alleles or by causing reproductive isolation itself through underdominance (Rieseberg 2001). We explored this in our dataset by asking if evidence of hybridization affected the number or size of inversions and found no support. That being said, our species comparisons are not necessarily sister species and do not necessarily have any geographic overlap, so they are not appropriate species to test models of speciation.

The location of inversions supports the hypothesis that they are often deleterious. We found that inversions were depleted for coding sequence and enriched for TEs consistently among the 32 genera (Figure 4). While we cannot eliminate the possibility that this reflects underlying biases in the inversion mutation rate or the reduced recombination effect after-the-fact, we think it is more likely to represent selection against inversions that disrupt genes or gene expression. Despite inversions often occurring in genes, the relatively small number of inversions compared to genes means that < 1 % of genes were affected by inversions in our dataset (Figure 5).

### The mechanism of inversions

We found a relatively small proportion of inversions contained duplications near inversion breakpoints (Figure 6). Taken at face value, this suggests non-homologous end-joining created a vast majority of inversions, but we cannot rule out error in the determination of inversion breakpoints. The current analysis uses one-to-one mappings to detect inversion regions, so if an inversion breakpoint were within a highly repetitive region, we may have underestimated the inversion size. Additionally, genome assemblies struggle resolving highly repetitive regions, such as centromeres (Naish et al. 2021). Although the genomes we included achieved chromosome-scale scaffolding, centromeric regions are especially dynamic and repetitive which means they are highly challenging to assemble correctly. Inversions can overlap with centromeres, and if breakpoints fell within centromeric regions we may not accurately locate them (Kirubakaran et al. 2020; Harringmeyer and Hoekstra 2022; Huang et al. 2022).

Depending on the mechanism, inversion mutations either create tandem duplicates at their breakpoints, or require them to be already present. With our current analysis, we cannot tell which is happening for each inversion, but future studies will be able as more chromosome-level genomes are produced. With multiple species genomes for a single genus, it will be possible to determine the derived and ancestral state for inversions and answer whether tandem duplications are a cause or consequence of inversions. Most current methods of genome comparison focus on pairwise comparisons (e.g., SyRI) but promising new methods using genomes graphs should allow for evolutionary analyses of genome structure across multiple species (Garrison and Guarracino 2022).

## Conclusion

Our results reject clock-like fixation of inversions in plants, and support early theoretical work that emphasized the central importance of selective and demographic effects on inversion fixation rates (Lande 1984). What we don’t know is what proportion of the inversions fixed were under positive selection versus weak negative selection. Future projects should combine population level sampling with comparative genomics to determine how often newly fixed inversions show signs of selection.

## Supplementary files

Supplementary Table 1: Species chosen for analysis and information collected, with full reference list for all genome assemblies included in the study.

Supplementary Figure 1: Distribution of inverted (blue) and syntenic (red) regions identified by SyRI in 32 species pair. The density of the sequence similarity score is plotted according to the corresponding nucleotide sequence identity where 1.00 = 100% identical.

Supplementary Figure 2: a) syntenic region sequence identity score distribution with vertical line representing the mean, b) inverted region sequence identity score distribution for the studied 32 species pair.

Supplementary Figure 3: Sequence divergence does not explain the proportion of the genome in inversions (*p*=0.506).

Supplementary Figure 4: The number of inversions, or proportion of the genome inverted for factors that may influence the accumulation of inversions (n=32). Hybridization (red), reproductive strategy (yellow), assembly methods parameters (blue).

