## Supplementary Figure for "The rate of inversion fixation in plant genomes is highly variable"

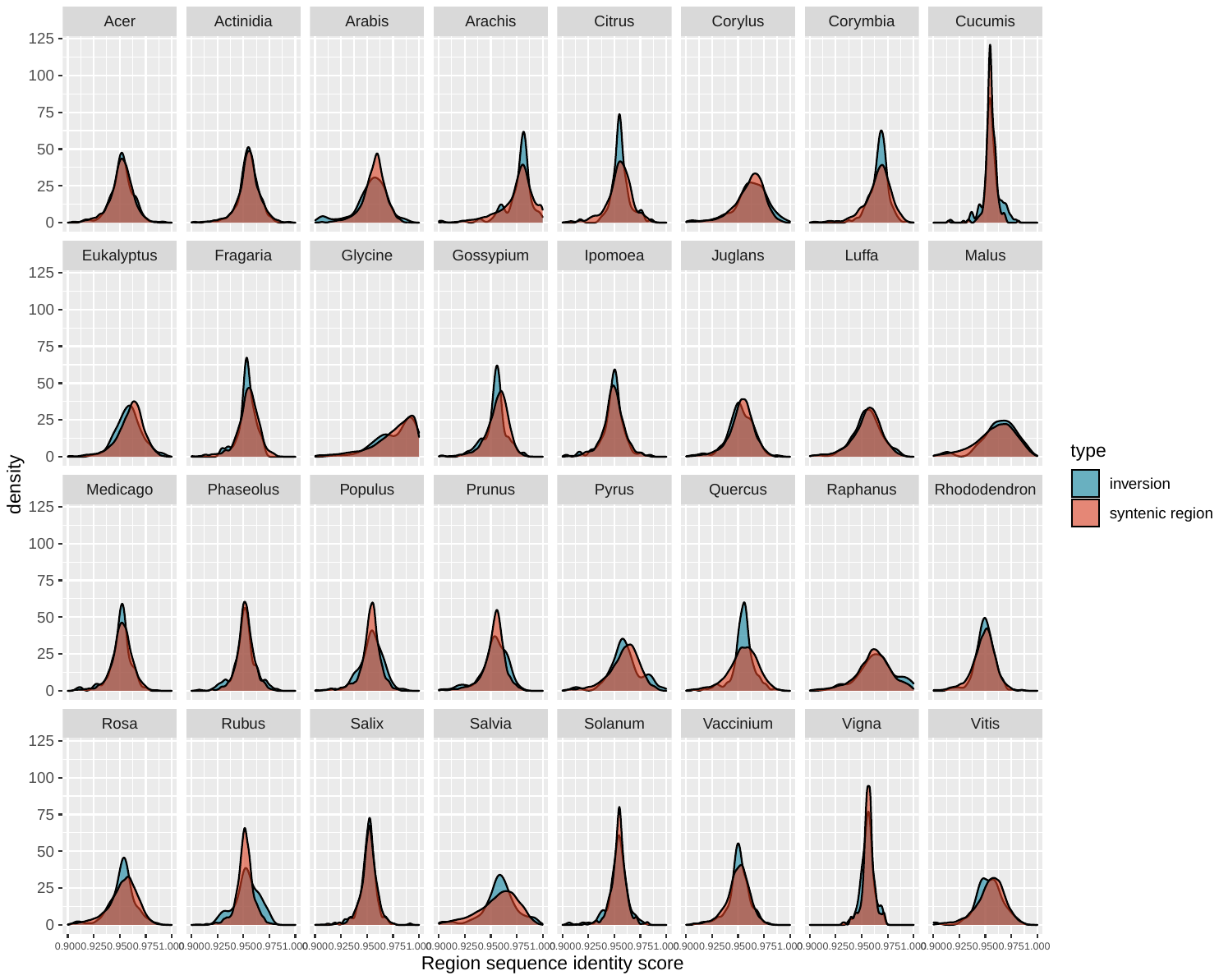

**Supplementary Figure 1**: Distribution of inverted (blue) and syntenic (red) regions identified by SyRI in 32 species pair. The density of the sequence similarity score is plotted according to the corresponding nucleotide sequence identity where 1.00 = 100% identical.

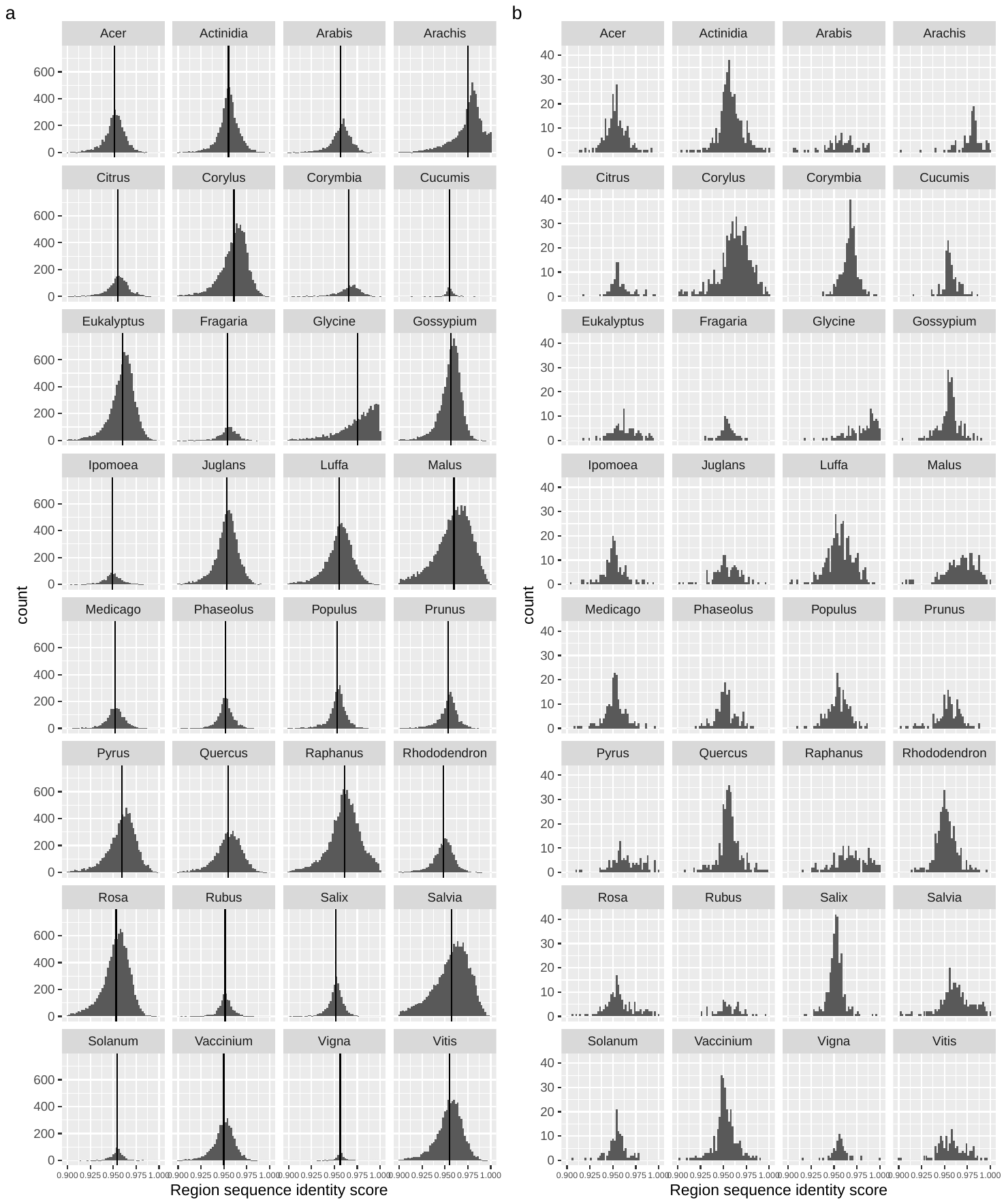

**Supplementary Figure 2**: a) syntenic region sequence identity score distribution with vertical line representing the mean, b) inverted region sequence identity score distribution for the studied 32 species pair.

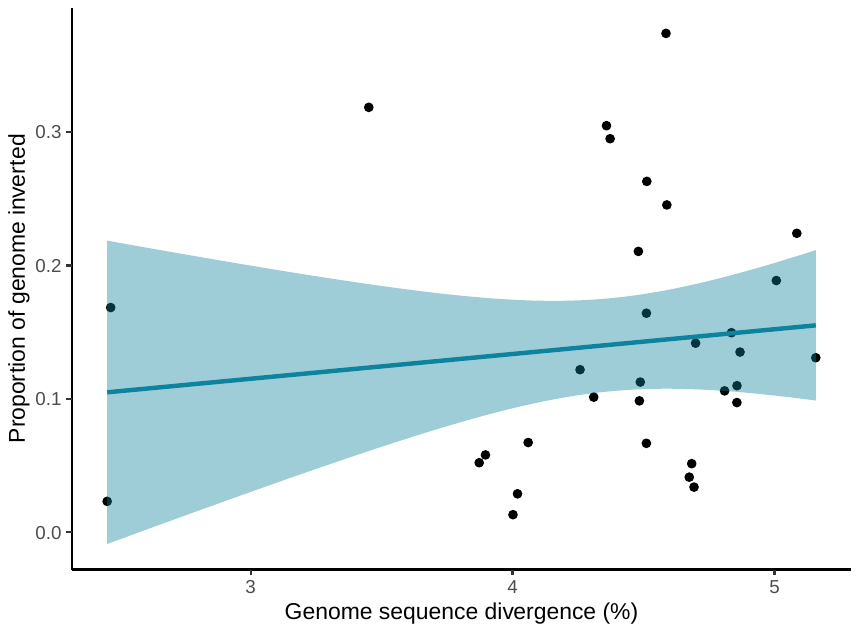

**Supplementary Figure 3**: Sequence divergence does not explain the proportion of the genome in inversions (*p*=0.506).

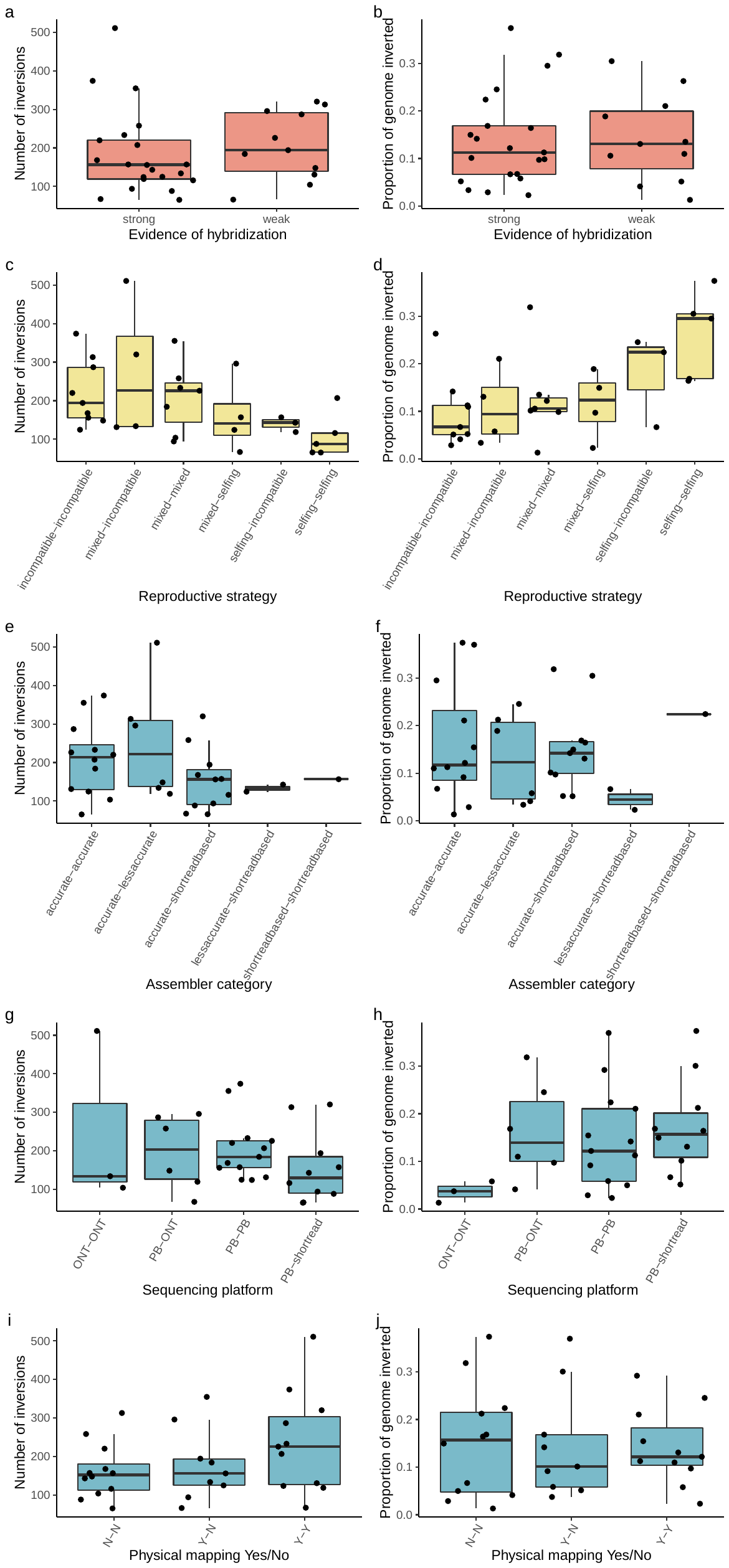

**Supplementary Figure 4**: The number of inversions, or proportion of the genome inverted for factors that may influence the accumulation of inversions (n=32). Hybridization (red), reproductive strategy (yellow), assembly methods parameters (blue).
